# *Fgf4* is critical for maintaining *Hes7* levels and Notch oscillations in the somite segmentation clock

**DOI:** 10.1101/2020.02.12.945931

**Authors:** Matthew J. Anderson, Valentin Magidson, Ryoichiro Kageyama, Mark Lewandoski

## Abstract

During vertebrate development, the presomitic mesoderm (PSM) is periodically segmented into somites, which will form the segmented vertebral column and associated muscle, connective tissue, and dermis. The periodicity of somitogenesis is regulated by a segmentation clock of oscillating Notch activity. Here, we examined mouse mutants lacking only *Fgf4* or *Fgf8*, which we previously demonstrated act redundantly to prevent PSM differentiation. *Fgf8* is not required for somitogenesis, but *Fgf4* mutants display a range of vertebral defects. We analyzed *Fgf4* mutants by quantifying mRNAs fluorescently labeled by hybridization chain reaction within Imaris-based volumetric tissue subsets. These data indicate that FGF4 controls Notch pathway oscillations through the transcriptional repressor, HES7. This hypothesis is supported by demonstrating a genetic synergy between *Hes7* and *Fgf4*, but not with *Fgf8*. Thus, *Fgf4* is an essential Notch oscillation regulator and potentially important in a spectrum of human Segmentation Defects of the Vertebrae caused by defective Notch oscillations.

## Introduction

A common developmental mode employed by many embryos is a segmentation clock that oscillates within a posterior growth zone; with each cycle, a segment forms. This stratagem has evolved independently within the three major bilaterian clades (annelids, arthropods, and chordates) as well as in plants ^1–5^. In chordates, a segmentation clock oscillates in the presomitic mesoderm (PSM); with each cycle, a pair of somites form flanking the neural tube ^6, 7^. Somites differentiate into dermis, skeletal muscle, tendons, as well as the vertebral column, which retains the segmented attribute of the somites ^8^.

In vertebrates, genes with an oscillatory pattern within the PSM include those encoding components or targets of the FGF, WNT and Notch signaling pathways ^7, 9, 10^. However, which individual genes oscillate differs among species, with the exception of the Notch-responsive HES/HER transcription factors, suggesting that Notch signaling is at the core of the somitogenesis clock ^9^. Supporting this idea, genetic and pharmacological manipulations demonstrate that Notch pathway oscillation in the mouse embryo is essential for somitogenesis^11^. In the mouse, oscillatory waves of the *Notch1* receptor and Delta-like 1 (*Dll1*) ligand expression, as well as the activated Notch receptor (cleaved Notch intracellular domain, NICD) sweep from the posterior to anterior, where oscillations arrest ^12, 13^. These oscillations are established through negative-feedback loops of Notch components such as the transcriptional repressor, HES7 ^14^, and the glycosyltransferase, LFNG ^15^, both of which are encoded by oscillating genes under Notch pathway regulation ^14^. Notch oscillations arrest in the anterior PSM, where NICD cooperates with TBX6 to periodically activate *Mesp2* expression ^16^. MESP2 then regulates formation of the nascent somite as well as its rostro-caudal patterning, which is essential for normal patterning of the subsequent vertebral column ^17, 18^.

Mutations in nearly all of the aforementioned Notch pathway genes have been identified in human patients where defective somitogenesis is thought to be the cause of Segmentation Defects of the Vertebrae (SDV) ^19, 20^. For example, frequently recessive mutations in *DLL3*, *HES7*, *MESP2* and *LFNG* and a dominant mutation in *TBX6* ^21, 22^ have been identified in patients with spondylocostal dysostosis (SCDO), which is characterized by severe vertebral malformations that include hemivertebrae, vertebral loss and fusion along the length of the axis^23^. Whereas SCDO is relatively rare, congenital scoliosis (CS), defined as a lateral curvature of the spine exceeding 10%, is much more common, with a frequency of 1:1000, which is suspected to be an underestimation because asymptomatic individuals do not seek medical care ^24^. Mutated alleles of *HES7*, *LFNG*, *MESP2*, and *TBX6* are all associated with CS ^22–25^.

The position in the anterior PSM where Notch oscillations arrest and *Mesp2* expression establishes the future somite boundary is called the determination front ^15, 18, 26^. This region is also the anterior limit of the wavefront in the classical clock-and-wavefront, a theoretical model proposed over 40 years ago to explain the precipitous and periodic formation of segments in the PSM ^27^. In this model, wavefront activity prevents the PSM from responding to the segmentation clock; hence somites form only anteriorly, where the wavefront ends. In chick or zebrafish embryos, exogenously added FGF protein or pharmacological inhibition of FGF signaling will shift the determination front rostrally or caudally, respectively. In the mouse, Cre-mediated inactivation of *Fgf4* and *Fgf8* specifically in the PSM results in an initial expansion of *Mesp2* expression, followed by the premature expression of somite markers throughout the PSM ^28^. Canonical WNT signaling is also a wavefront candidate, mostly because ectopic activation of βcatenin in the PSM results in an expansion of this tissue and a rostral shift of *Mesp2* expression ^29, 30^. However, this expansion observed in βcatenin “gain-of-function” mutants requires FGF4 and FGF8 activity, suggesting that these FGF signals are synonymous with the wavefront ^28^.

In addition to its role in preventing somite differentiation, FGF signaling is also implicated in the regulation of key Notch components of the segmentation clock. Several studies have demonstrated that pharmacological inhibition of FGF disrupts Notch oscillations ^31, 32^. In mouse embryos with PSM-specific loss of *FGF receptor 1* or both *Fgf4* and *Fgf8*, Notch pathway genes such as *Hes7*, are downregulated, although this analysis can be complicated due to the loss of PSM tissue in these mutants ^28, 32^. Here, we focus on FGF-Notch interactions by analyzing mouse mutants that lack only one of the wavefront *Fgf* genes, *Fgf8* or *Fgf4*. An indispensable technique in our analysis is whole mount *in situ* hybridization chain reaction (HCR), which allows us to multiplex different gene expression domains and quantify mRNA levels within specific embryonic tissues ^33–35^. We demonstrate that, while *Fgf8* is not required for somitogenesis, *Fgf4* is required for normal Notch oscillations and patterning of the vertebral column.

## Results

### FGF4 activity is required in the presomitic mesoderm for normal segmentation of rostral somites

To analyze the role of *Fgf4* or *Fgf8* expression in the primitive streak and PSM (Figure 1A - B), we inactivated each gene specifically within these tissues using TCre transgenic activity ^36^, thus generating “*Fgf4* mutants” (TCre; *Fgf4* ^flox/Δ^; “Δ”= “deleted” or null) or “*Fgf8* mutants” (TCre; *Fgf8* ^flox/Δ^; see Table 1). Whereas *Fgf8* mutants do not survive much beyond birth due to kidney agenesis ^36^, *Fgf4* mutants are viable and found at Mendelian ratios at weaning (n = 18 controls and n = 21 mutants). Skeleton preparations of *Fgf8* mutant embryos at E18.5 (n = 22) revealed all vertebral bodies were present and normally patterned. *Fgf8* mutants also presented with minor cervical and/or lumbar homeotic transformations, each with incomplete penetrance and expressivity: small ribs were sometimes present in the most posterior cervical vertebra (8/22, 4 bilateral and 4 unilateral) or on the most anterior lumbar vertebra (8/22, 4 bilateral and 4 unilateral). On the other hand, *Fgf4* mutants display a variety of segmentation defects in the cervical and thoracic vertebrae with 100% penetrance (Compare Figure 1D, D’ with C). An average of 7.9 defects occurred per mutant and consisted of hemivertebrae, and misshapen and deleted vertebrae (Figure 1E).

**Figure 1.**
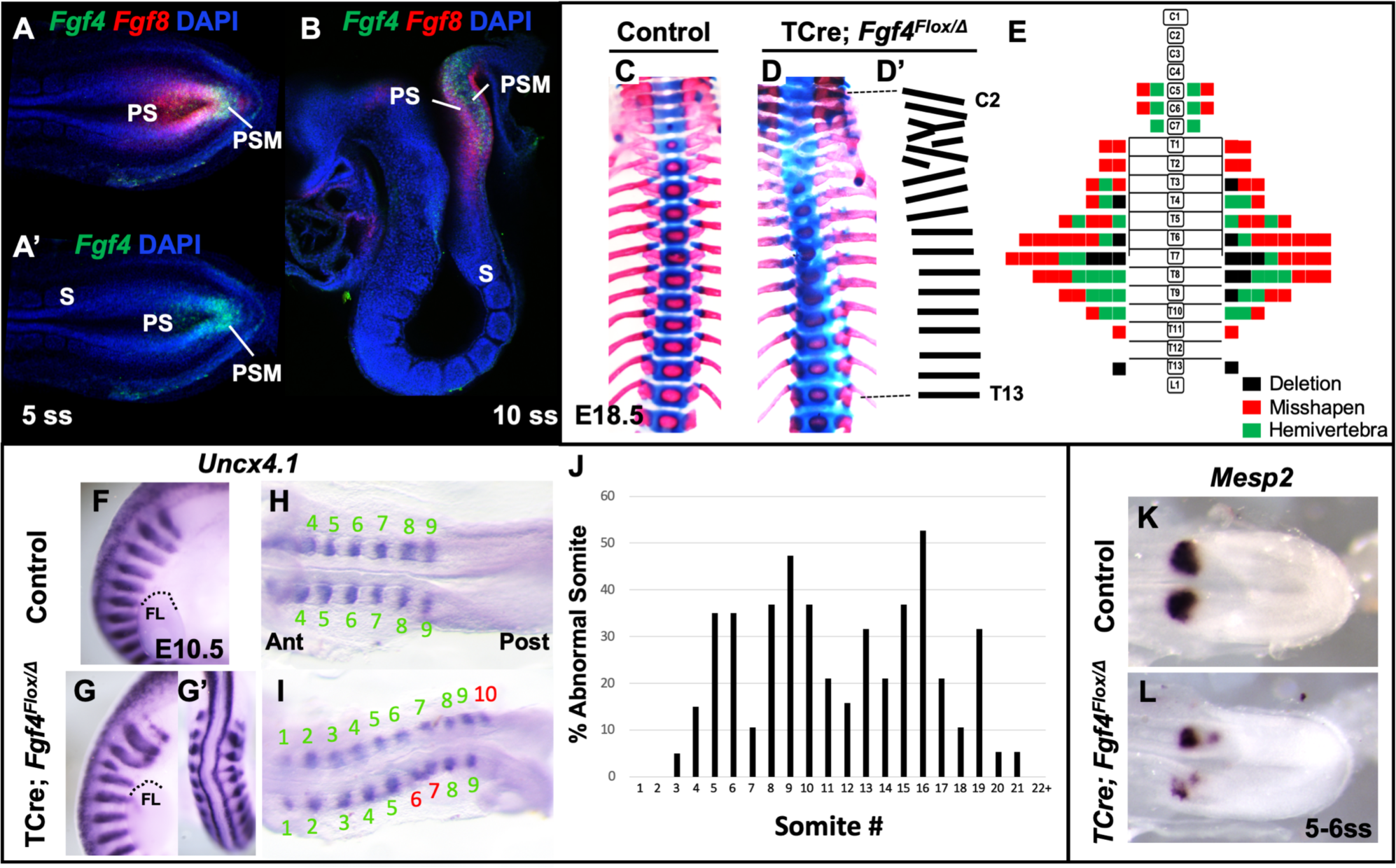
*Fgf4* mutants have defects in vertebral and somite patterning. **A, A’)** Max intensity projection (MIP) of HCR staining of *Fgf4* and *Fgf8* mRNA expression (A) or *Fgf4* only (A’) in the PSM of a 5 somite stage wildtype embryo (dorsal view). PS: primitive streak, PSM: presomitic mesoderm, S: somite. **B)** MIP of 4 parasagittal z-sections (approximately 5um each) of HCR staining of *Fgf4* and *Fgf8* mRNA expression, of a 10 somite stage wildtype embryo (lateral view, mediolateral to the midline). **C, D**) Skeletal preparations (ventral view after removing ribcage) of E18.5 control and *Fgf4* mutants. **D’)** Tracing of the *Fgf4* mutant vertebral pattern in D. **E)** Histograms representing the vertebral column and associated ribs (C = cervical, T = thoracic, L = lumbar), showing variety and location of vertebral defects in E18.5 *Fgf4* mutants (n = 12). Each block in the histogram represents a single defect as indicated in the key, bottom right. **F-I)** Wholemount *in situ* hybridization (WISH) detection of *Uncx4.1* mRNA expression in control and *Fgf4* mutants at E10.5 (F, G lateral views, anterior top; G’ dorsal view of embryo in G) and at 9-10 somite stage (H, I dorsal view, anterior left); dotted line in F and G marks the anterior boundary of the forelimb (FL). **J)** Graph depicting the percentage of abnormal somites scored with a mispatterned *Uncx4.1* mRNA WISH pattern (F-I) in *Fgf4* mutants at the somite position listed on the x-axis. Somites 1-6 were scored in somite stage 7 *Fgf4* mutants (n=20) and somites 7-26 were scored in 26-30 somite stage *Fgf4* mutants (n=19). Note that posterior to somite 21 the Uncx4.1 pattern is normal. **K**, **L)** Dorsal view (anterior left) of 5-6 somite stage control (K) and *Fgf4* mutant (L) embryos WISH-stained for *Mesp2* mRNA expression. In all control embryos (n=7) the *Mesp2* pattern was normal; in 5/8 mutant embryos, the pattern was abnormal.

**Table 1.**
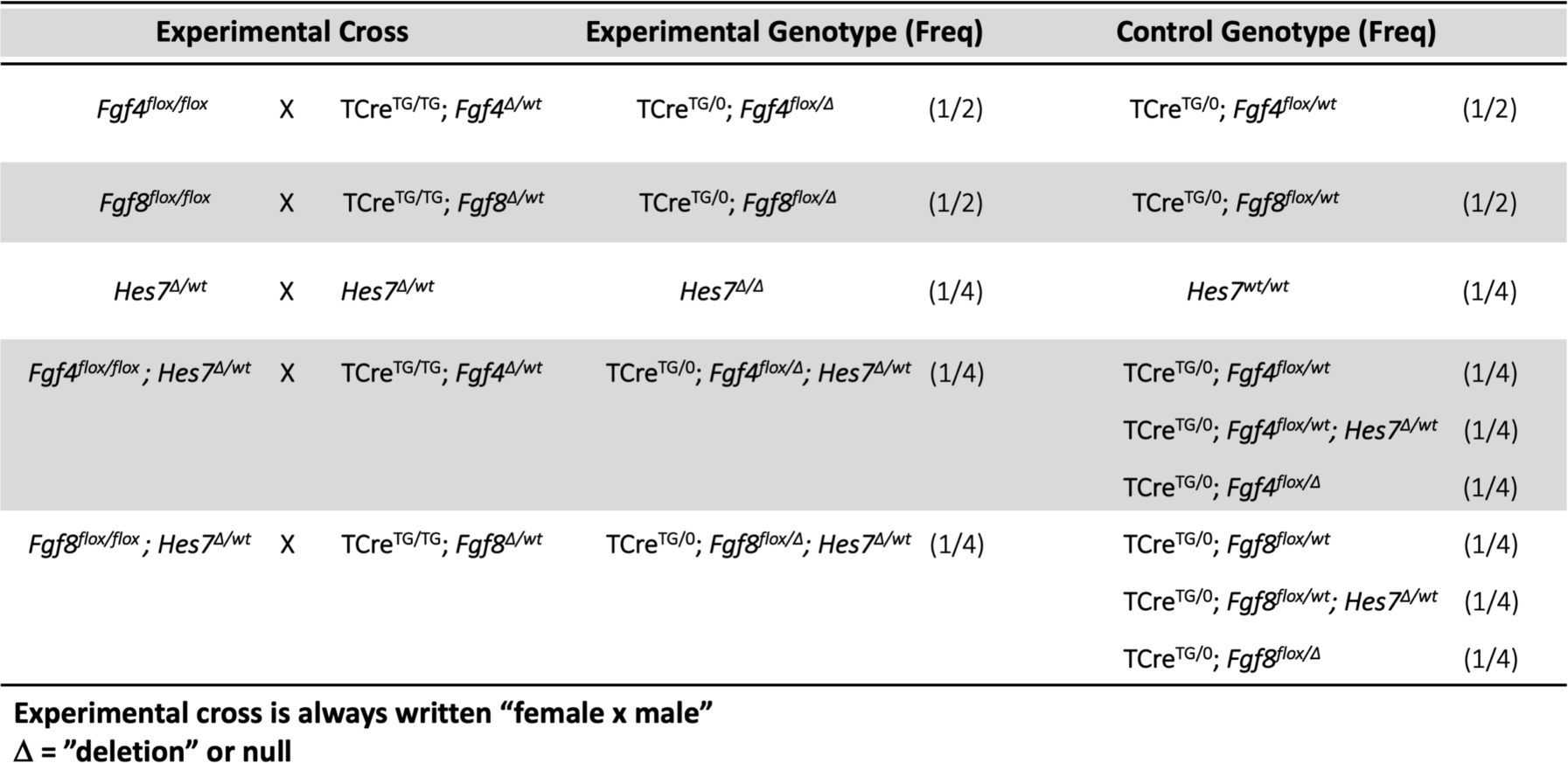
Experimental Crosses.

We examined gross somite patterning in *Fgf4* mutants by staining embryos at various stages for *Uncx4.1* mRNA, which marks the posterior somite compartment ^37, 38^. Analysis of 39 *Fgf4* mutants revealed a frequency of irregular *Uncx4.1* expression specifically in future cervical and thoracic somites with full penetrance (Figure 1F-J).

To determine if these malformed somites were due to an error in segmentation, we examined *Mesp2* expression, which occurs in the anterior presomitic mesoderm (PSM) and is required for normal somite segmentation and rostral-caudal somite identity ^18^. At the stages when irregular somites are emerging from *Fgf4* mutant PSM, we detected aberrant *Mesp2* expression, whether we analyzed gene expression by traditional wholemount *in situ* hybridization (WISH) staining (Figure 1K-L) or by fluorescent HCR analysis (Figure 2A, B). However, at later stages (22-24 somite stages), when properly segmented somites are emerging from *Fgf4* mutant PSM, *Mesp2* expression appears normal (Figure 1 - figure supplement 1). Therefore, we conclude that this indistinct pattern of *Mesp2* expression at early stages is likely responsible for the vertebral defects in *Fgf4* mutants.

**Figure 2.**
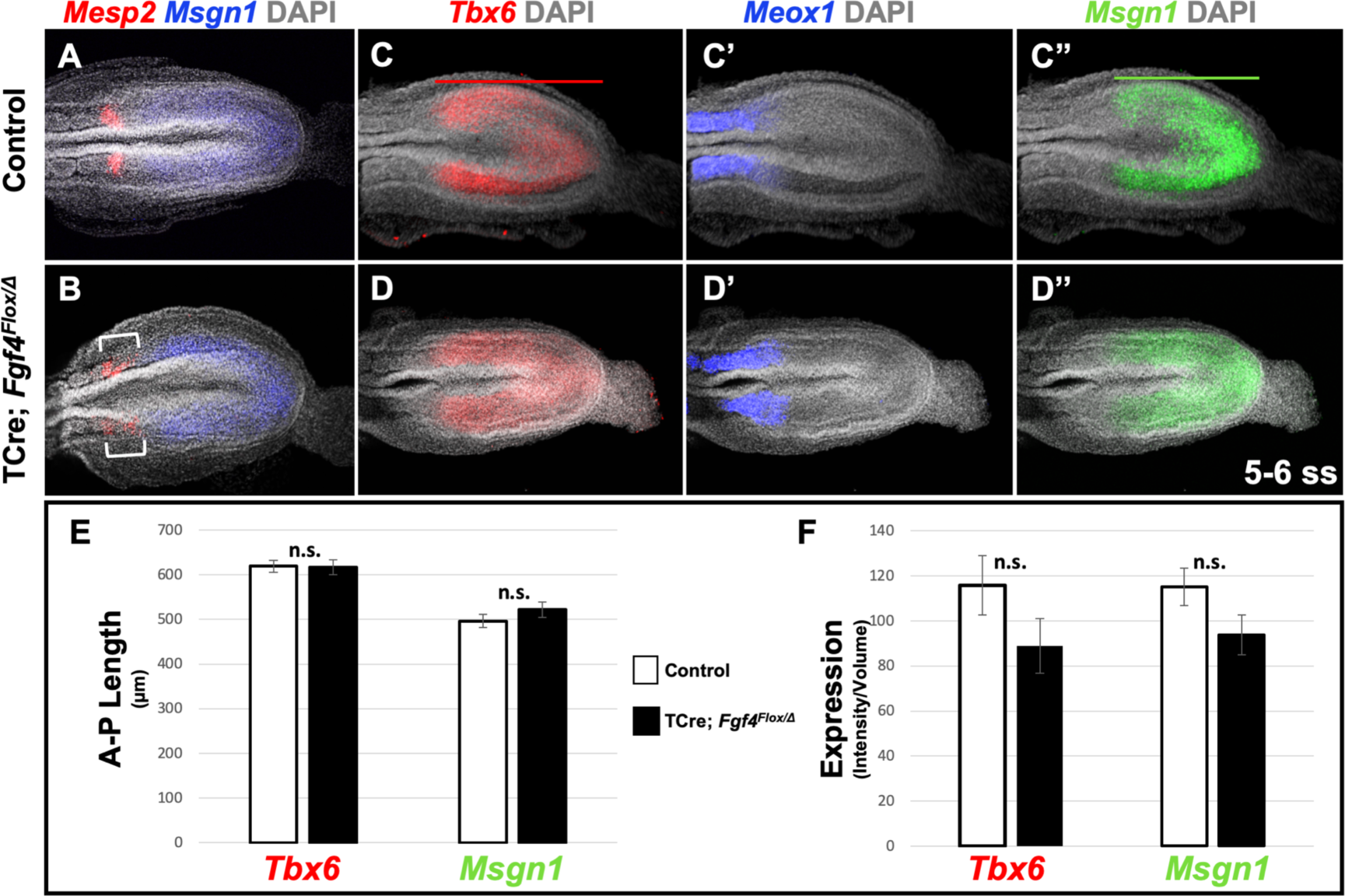
*Fgf4* mutants maintain a normal wavefront. **A**, **B)** HCR staining of 5-6 somite stage control (A, n = 8) and *Fgf4* mutant embryos (B, n = 11). In mutants, *Msgn1* expression was similar between mutants and controls, but *Mesp2* expression was abnormal (6/11). **C-D’’)** HCR staining of 5-6 somite stage control and *Fgf4* mutant embryos showing normal expression of *Meox1* (control, n = 4; mutant, n = 4), *Msgn1* (control, n = 9; mutant, n = 8), and *Tbx6* (control, n = 10; mutant, n = 8). **E)** There is no significant difference in the anterior-posterior length of *Tbx6* and *Msgn1* domains in control (n = 9) and mutant embryos (n = 8; green and red bars in C and C’). **F)** Quantification of *Tbx6* and *Msgn1* mRNA expression, determined by measurement of fluorescence intensity per cubic micrometer within the volume of the *Msgn1* or *Tbx6* domain of expression, respectively. There is no significant difference between control and mutant embryos. In E and F, data are mean±s.e.m, significance determined by a Student’s t-test. *Msgn1*: control n=7; mutant n = 8. *Tbx6*: control, n = 9; mutant, n = 8. All images: MIP, dorsal view, anterior left. Same embryo shown in C-C’’ and D-D’’.

### Wavefront gene expression is normal in *Fgf4* mutants

For the rest of our mRNA expression analysis we mostly relied on whole mount *in situ* HCR in conjunction with Ce3D++ tissue clearing. Because the fluorescent signal intensity corresponds proportionally to the number of mRNA molecules ^35^, HCR allows for quantitative and multiplexed mRNA detection ^34^. Accurate quantification of the HCR signal requires a method such as Ce3D++ tissue clearing (see Materials and Methods), which allows for imaging in deep tissues without signal loss.

We first addressed why a distinct and symmetrical band of *Mesp2* expression fails to form in *Fgf4* mutants. *Mesp2* expression is normally triggered by activated Notch (NICD) and TBX6 at the determination front in the anterior PSM, where cells fall below a critical threshold of wavefront FGF signaling ^16^. We previously showed that this FGF activity was encoded by *Fgf4* or *Fgf8;* if we inactivated both *Fgfs* with TCre (*Fgf4/Fgf8* double mutant), *Mesp2* expression is transiently activated throughout the PSM, which then aberrantly expresses the paraxial mesoderm marker, *Meox1* ^28^. To determine if the wavefront position is altered in *Fgf4* mutants we examined the anterior-posterior length and expression levels of genes responsive to this activity at early somite stages when aberrant *Mesp2* expression occurs. Both *Msgn1* and *Tbx6* are required to specify the paraxial mesoderm ^39–41^, and are silenced in the absence of wavefront activity in *Fgf4/Fgf8* double mutants ^28^. Neither the length nor level of expression of *Tbx6* and *Msgn1* were significantly altered in *Fgf4* mutants (Figure 2C, C”, D, D”, E, F). Consistent with this observation, the paraxial differentiation marker, *Meox1*, was not expanded into the PSM (Figure 2 C’ and D’), as occurs in *Fgf4/Fgf8* double mutants ^28^.

Therefore, we conclude that wavefront activity is normal in *Fgf4* mutants, an insight consistent with the observation that normal axis extension occurs in these mutants, with no loss of caudal vertebrae ^42^. We surmise that this normal wavefront position and signaling in *Fgf4* mutants are likely maintained by *Fgf8*, which is expressed at normal levels (Figure 3A-C) and is sufficient for maintaining normal expression levels of the FGF target genes, *Spry2*, *Etv4*, and *Spry4* (Figure 3D-F’’’); these FGF targets are silenced in *Fgf4/Fgf8* double mutants ^28^. Therefore, neither a change in wavefront activity nor a change in determination front position explains the aberrant pattern of *Mesp2* expression in *Fgf4* mutants.

**Figure 3.**
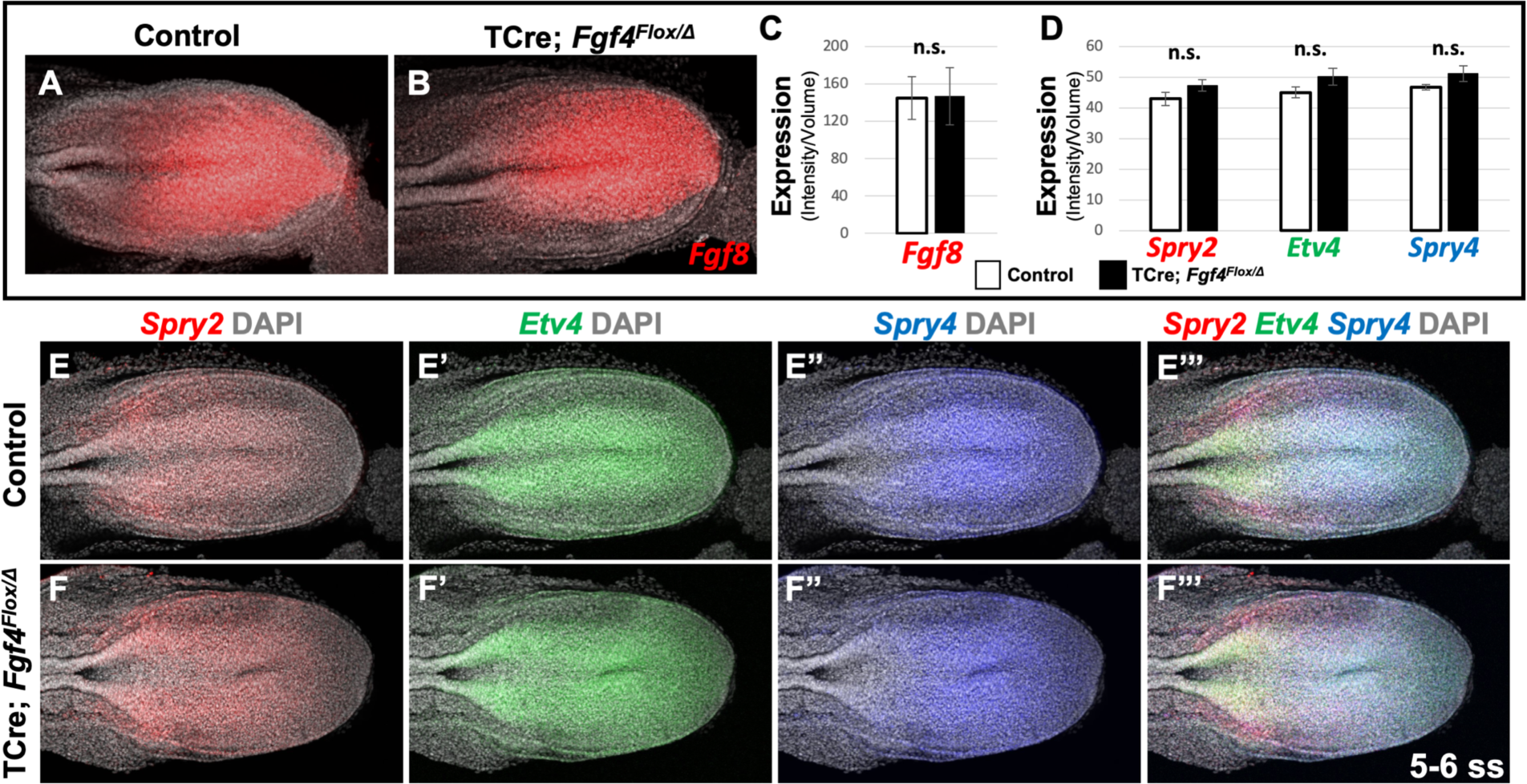
*Fgf8* is sufficient for maintaining canonical FGF-responsive gene expression in *Fgf4* mutants. **A**, **B)** HCR staining of 5-6 somite stage control (n=6) and *Fgf4* mutant (n=3) embryos showing comparable expression of *Fgf8,* quantified in C). **C)** Quantification of *Fgf8* mRNA expression, determined by measurement of fluorescent intensity per cubic micrometer within the volume the mesoderm-specific *Tbx6* expression domain. Note, there is no significant difference between control and mutant embryos. **D)** Quantification of HCR analysis of *Spry2*, *Etv4*, and *Spry4* expression in 5-6 somite stage control (n =4) and *Fgf4* mutant (n =4) embryos imaged in **E-F’’’**. Data in C and D are mean±s.e.m, significance determined by a Student’s t-test. All images: MIP, dorsal view, left. Same embryo shown in E-E’’’ and F-F’’’.

### Oscillation of Notch family components is altered in the PSM of *Fgf4* mutants

We then examined the pattern of activated Notch in *Fgf4* mutants by immunostaining for Notch intracellular domain (NICD), which is an obligate factor in the transcriptional complex that activates *Mesp2* ^17^. Overall NICD levels were unchanged between *Fgf4* mutants and littermate controls at the embryonic stages when somitogenesis was abnormal (Figure 4A). At these stages, we always observed two to three distinct stripes of NICD along the anterior-posterior axis in both mutant and littermate control PSM (Figure 4B-C), demonstrating the oscillatory nature of Notch activation ^12, 15, 43^. However, mutant oscillatory stripes of NICD were always less distinct, compared to controls. This was more evident when the relative intensities of NICD immunostaining signals were modeled using Imaris software. In controls, cells medial-lateral to each other had similar levels of activated NICD, whereas in mutants, this coordination was less distinct (Figure 4B’-C’). Importantly, this blurred pattern occurred in the anterior PSM, where NICD activates *Mesp2* expression at the determination front (Figure 4B’-C’, brackets).

**Figure 4.**
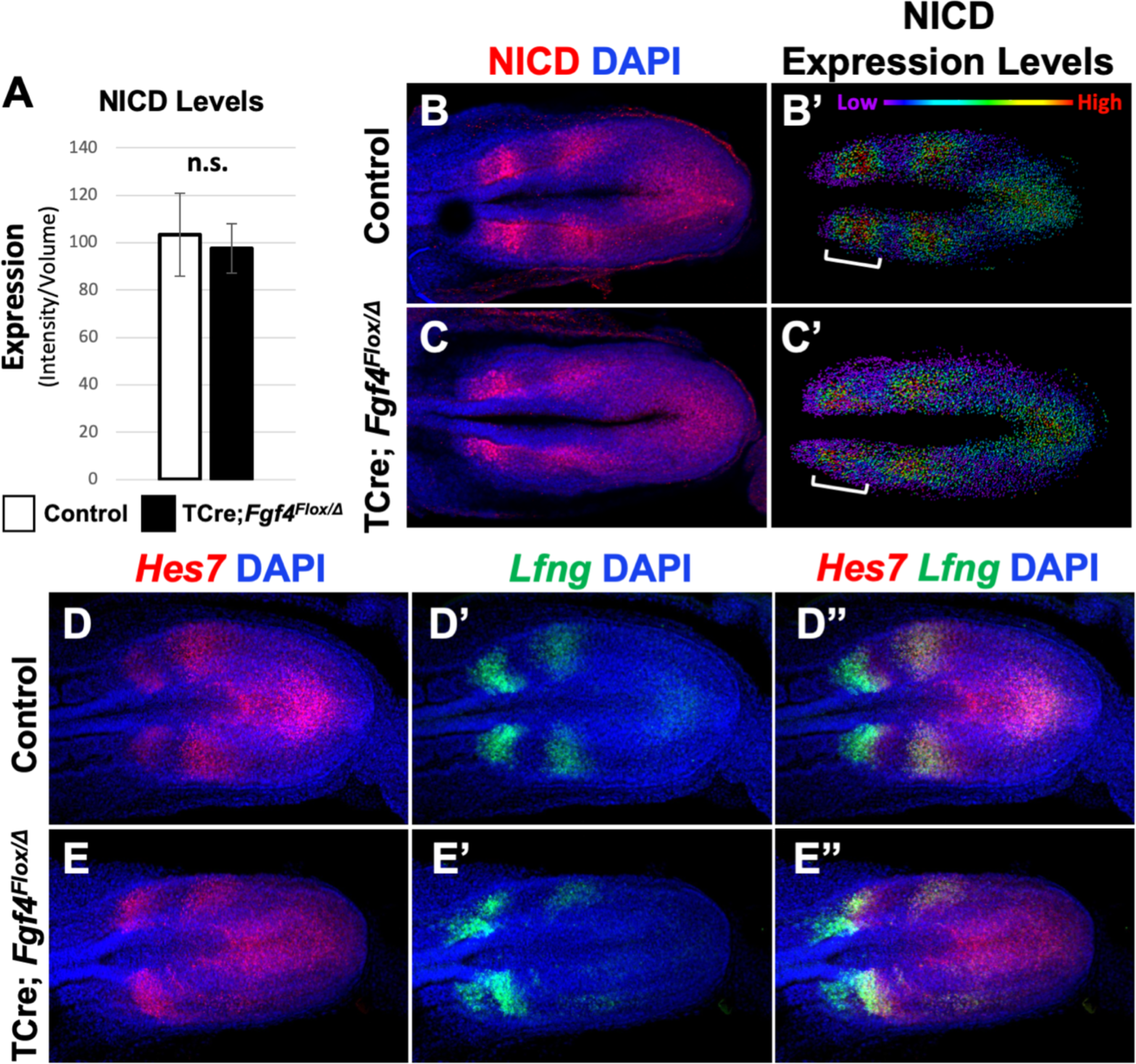
Pattern of Notch signaling is abnormal in *Fgf4* mutants. **A)** Quantification of immunostained NICD fluorescent signal specifically in the mesoderm of 5-6 somite stage embryos. Note, there is no significant difference between *Fgf4* mutants (n=5) and littermate controls (n=5). Data are mean±s.e.m, significance determined by a Student’s t-test. **B, C)** 5-6 somite stage control (B) or *Fgf4* mutant (C) embryos immunostained for NICD. **B’ C’)** Same images as in (B,C) but visualized using the Imaris spot modeling function where each fluorescent signal is represented as a colored sphere according to pixel intensity (lowest value is purple and highest value is red, as indicated). Note that the pattern in the *Fgf4* mutant (C’) is less distinct and blurred compared to littermate control (B’); brackets indicate anterior PSM. **D-E’’)** HCR staining of representative 5-6 somite stage control (n = 10) and *Fgf4* mutant (n = 8) embryos for the indicated genes. Note *Fgf4* mutant expression of *Hes7* and *Lfng* in mutants is abnormal. All images: MIPs, dorsal view, anterior left. Same embryo shown in B, B’; C, C’; D-D’’; E-E’’.

A normal NICD pattern is achieved through a number of negative feedback loops including the transcriptional repressor, HES7 ^14^ and the glycosyltransferase, LFNG ^15^; disruption of either gene encoding these factors causes uncoordinated Notch signaling ^14^. *Hes7* and *Lfng* are themselves targets of activated Notch and therefore their expression oscillates ^14^. Although oscillatory patterns of *Hes7* and *Lfng* are apparent in both *Fgf4* mutants and littermate controls (Figure 4D- E’’), the mutant patterns are clearly uncoordinated and indistinct, similar to the NICD immunostained mutant (Figure 4D-E’’). Together, these data suggest the aberrant *Mesp2* pattern in *Fgf4* mutants can be explained by imprecise oscillations of Notch signaling.

### *Hes7* expression is reduced in the PSM of *Fgf4* mutants

We proceeded to analyze the expression of *Hes7* in greater detail in *Fgf4* mutants. *Hes7* is within the *Hes/Her* class of transcriptional repressors that are the only oscillating clock genes conserved amongst mouse, chicken, and zebrafish ^9^. In the mouse, mutations that accelerate HES7 production will accelerate the tempo of the segmentation clock ^44^. Hence, *Hes7* is considered to be a fundamental pacemaker of the segmentation clock that controls somitogenesis ^45^.

Analysis of *Hes7* mRNA in *Fgf4* mutants at 5-6 somite stages, using conventional WISH, revealed a reduced and aberrant expression pattern in *Fgf4* mutants (Figure 5-figure supplement 1). Analysis with HCR was much more informative, allowing unambiguous pattern analysis (Figure 4) and quantification (Figure 5, Figure 5-figure supplement 2, Figure 5-figure supplement 3). Characterization of an oscillatory gene expression pattern in the PSM is achieved by classifying static expression patterns into three phases ^46^. This approach has been used to characterize *Hes7* expression at E9.5-E10.5, when one to two stripes of expression are observed ^47, 48^. However, in the E8.5 PSM, when aberrant Notch signaling occurs in *Fgf4* mutants, there are two to three *Hes7* expression stripes in both controls and *Fgf4* mutants, indicating a faster somitogenesis clock at this stage (Figure 4, 5). Therefore, we generated criteria for classification of phases at E8.5 as follows. Phase I: three stripes with the most posterior stripe limited to the posterior midline and the most anterior stripe having reached the anterior boundary of the PSM (Figure 5A). Phase II: two stripes with the posterior stripe having expanded laterally compared to phase I and the anterior stripe having not reached the anterior limit of the PSM (Figure 5B).

**Figure 5.**
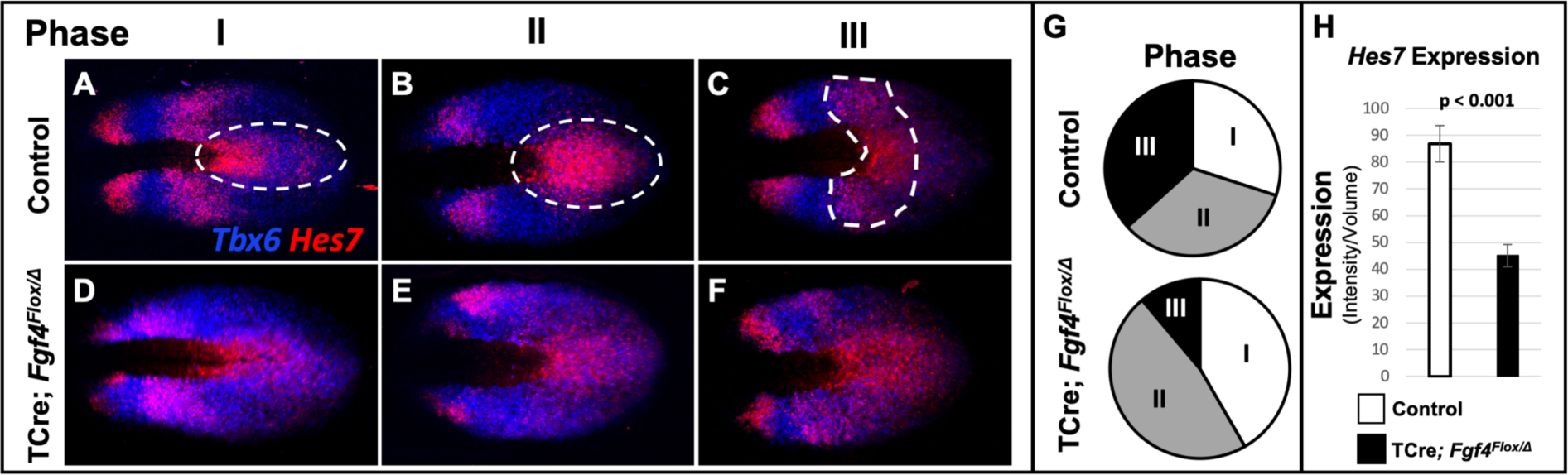
*Hes7* expression is abnormally patterned and reduced in *Fgf4* mutants. **A-F)** HCR staining for *Hes7* expression in 5-6 somite stage control and *Fgf4* mutant embryos sorted by phase of oscillation, showing abnormal expression of *Hes7* in mutants. **G)** Distribution of phases in control and *Fgf4* mutants shown in A-F; controls: phase I, n = 9; phase II, n = 10; phase III, n = 11. *Fgf4* mutants: phase I, n = 6; phase II, n = 9; phase III, n = 2. **H)** Quantification of *Hes7* expression specifically within the PSM (all phases combined) in 5-6 somite stage control (n=11) and *Fgf4* mutant (n=8) embryos shows significantly decreased expression in the *Fgf4* mutant. Quantification of *Hes7* mRNA expression, determined by measurement of fluorescence intensity per cubic micrometer within the volume the mesoderm-specific *Tbx6* expression domain. Data in H are mean±s.e.m, significance determined by a Student’s t-test. All images are MIPs, dorsal view, anterior to left.

Phase III: two distinct stripes and a third stripe initiating at the posterior midline (Figure 5C). Control embryos are distributed nearly equally between each phase (Figure 5G) as is the case for *Hes7* expression in older embryos ^47, 48^. It is challenging to place *Fgf4* mutants in phases, emphasizing the aberrant *Hes7* expression pattern. However, in our assessment, we allocated *Fgf4* mutants within all three phases (Figure 5D-F), with most found in oscillation phase II (Figure 5G).

*Hes7* is expressed in both the PSM and neural ectoderm. To quantify expression only in the PSM, we used Imaris software to create a volumetric model of the PSM, based on *Tbx6* expression, which is limited to the PSM at these stages ^49^. By measuring the intensity of the *Hes7* HCR signal within this volume, we obtained a PSM-specific *Hes7* quantification for each embryo (see Figure 5- figure supplement 3 and Material and Methods). PSM-specific *Hes7* expression is reduced in *Fgf4* mutants to 51.8 % of that expressed in litter mate controls (Figure 5H). A comparison between mutant and control *Hes7* expression levels within each phase reveals that the reduction in *Fgf4* mutants is not due to the difference in phase allocation between *Fgf4* mutants and littermate controls (Figure 5 - figure supplement 4). Moreover, after the 22 somite stage, when forming somites are normally segmented in mutants (Figure1J), there is no significant difference in *Hes7* PSM expression levels between *Fgf4* mutants and controls (Figure 5 - figure supplement 5). Therefore, we conclude that a reduction of *Hes7* expression may account for the defective segmentation during rostral somitogenesis in *Fgf4* mutants.

To correlate *Hes7* expression levels with pattern, we created Imaris-generated spot models of the HCR signal, which correspond to a single mRNA molecule or clusters of mRNA molecules within a subcellular-volume (see Materials and Methods). We then colored-coded individual spots to reflect the intensity of the *Hes7* HCR signal, using a linear series of intensity-based cutoffs every 20% to generate five colors (quintiles) (Figure 6A-B). In controls, the lowest intensity quintile (blue) contains the most spots (40%) and each higher intensity group contains 10% fewer spots (Figure 6A, white bars in C). The distribution of spots in the *Fgf4* mutant, compared to controls, is significantly skewed towards the lowest quintile at the cost of higher-level expression spots (Figure 6B, black bars in C). This modeling provides an effective illustration of the pattern of *Hes7* expression.

**Figure 6.**
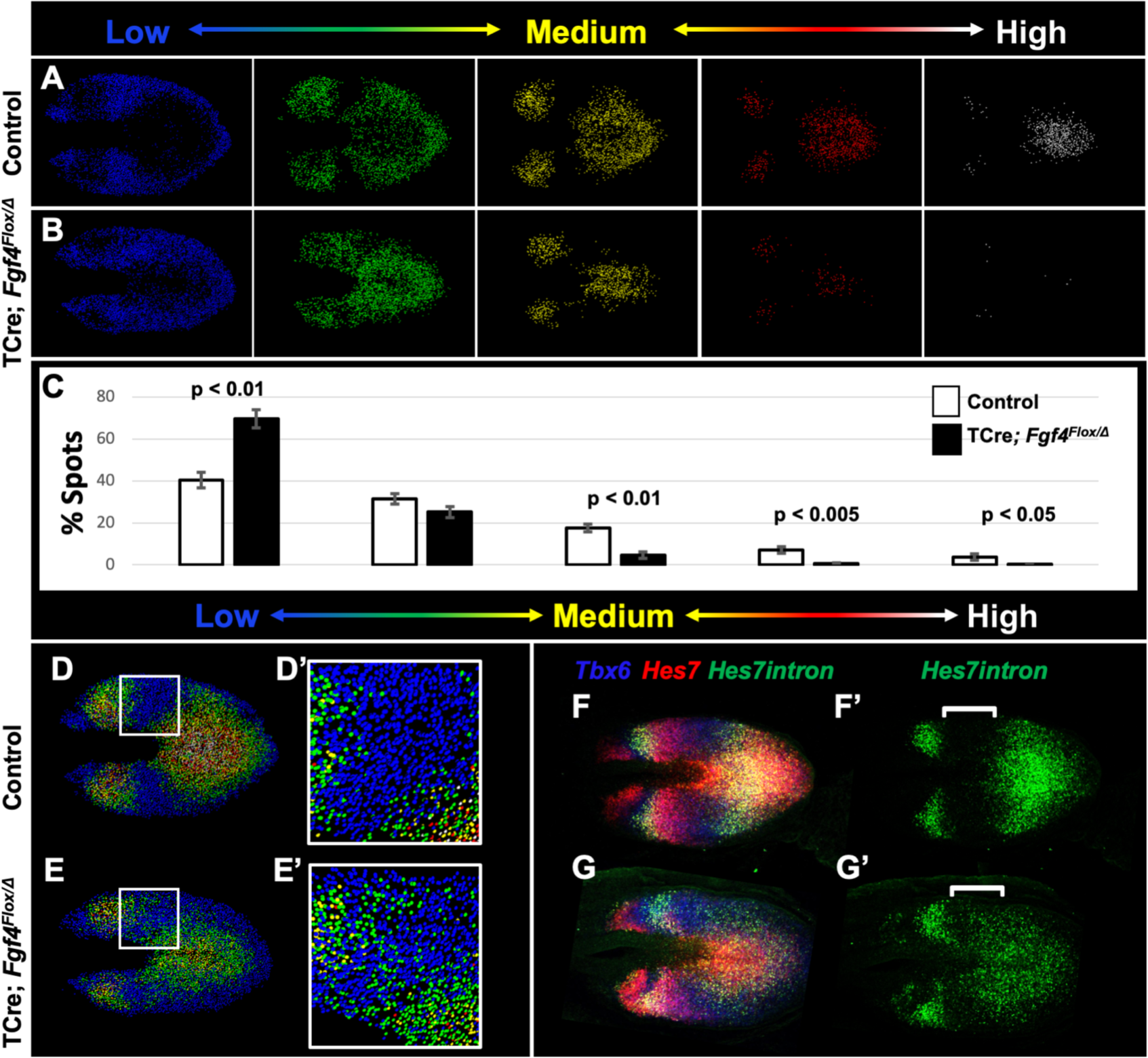
Reduced *Hes7* expression with less distinct peaks and troughs in the *Fgf4* mutant PSM. **A, B)** Spot models based on HCR analysis of *Hes7* expression, in 5-6 somite stage control and *Fgf4* mutant embryos within the PSM, as defined by the *Tbx6* expression domain. Localized *Hes7* expression is colored by level of expression from low (blue) to high expression (white), as indicated. Note there are less high expressors (yellow, red and white) and more low expressors (blue). This insight is quantified in **C)**, where the percentage of spots found in each expression-level group is graphed. Data are mean±s.e.m, significance determined by a Student’s t-test; control, n = 6; mutant, n = 5. **D-E’)** Composite of spot models from A and B. Note that the *Hes7* expression trough (boxed region, expanded in D’ and E’) between anterior and posterior oscillatory peaks is less distinct in mutant (E’) with more higher expressors (green) than in control embryos (D’). **F-G’)** MIP of HCR staining of 5-6 somite stage control and *Fgf4* mutant embryos using probes against *Tbx6*, *Hes7*, and *Hes7intron* which specifically labels active *Hes7* transcription. Note the trough of *Hes7intron* signal between peaks (brackets) in mutants is less pronounced than controls, indicating a failure of transcriptional repression of *Hes7* in this region. Representative embryos are shown; control, n = 6; mutant, n = 5. All images are dorsal views, anterior left. Same embryo shown in F, F’; G, G’.

In control embryos, each stripe of *Hes7* expression contains a concentric gradient of signal intensity with highest expression at the center (white spots) and lowest expression at the outside (blue spots) (Figure 6A, D). *Fgf4* mutants maintain some of this concentric organization, but the boundaries between groups is less clear (Figures 6B, E); in particular the trough between peaks of *Hes7* expression is more shallow, and contains more higher-intensity spots (mostly second quintile, green) than littermate controls (Figures 6D’ and E’). To determine if this indicated a failure to repress *Hes7* transcription in these regions, we performed HCR using a probe that hybridized to the *Hes7* intronic sequences, and therefore specific to newly transcribed pre-mRNA. This analysis revealed that transcription of *Hes7* is reduced in the posterior PSM and is more widespread, extending into the trough between *Hes7* mRNA peaks in the *Fgf4* mutant (white brackets in Figure 6 F’, G’). Therefore, we hypothesized that FGF4 is required to maintain *Hes7* transcription in the PSM above a threshold required for normal Notch oscillation. The reduced level of HES7 in *Fgf4* mutants is insufficient to fully autorepress its own expression, resulting in an aberrant pattern of expression.

### A synergistic defect in *Fgf4/Hes7* mutants reveals that *Fgf4* is required to maintain *Hes7* above a critical threshold

To test our hypothesis that a reduction of *Hes7* expression causes the *Fgf4* mutant vertebral defects, we asked if these defects worsen if we further reduce *Hes7* expression by removing one gene copy. We compared such mutant to littermate controls that were simple *Fgf4* mutants (with two wildtype *Hes7* alleles) or *Hes7* heterozygotes. These *Hes7* heterozygotes also carried TCre and one floxed *Fgf4* allele (see Table 1, and Figure 7) resulting in *Fgf4* heterozygosity in the TCre expression domain. However, *Fgf4* heterozygosity had no effect on the phenotype because the defects we observed were similar to the *Hes7* heterozygous defects reported by the Dunwoodie lab ^25^. About 50% of our *Hes7* heterozygotes had defective lower thoracic vertebrae with a frequency of 2.8 defects per animal. In littermate *Fgf4* mutants, vertebral defects were completely penetrant, with an average of 7.7 defects per animal, a frequency similar to that of progeny in our original genetic cross (Figure 1). However, compound *Fgf4/Hes7* mutants displayed a large set of defects with 25 defects per animal (Figure 7C-E), a frequency significantly greater than littermate *Fgf4* mutants (3-fold greater, p < 0.005) or *Hes7* heterozygotes (9-fold greater, p < 0.001). These data indicate a synergistic, as opposed to additive, effect of loss of one *Hes7* allele in the *Fgf4/Hes7* compound mutant.

**Figure 7.**
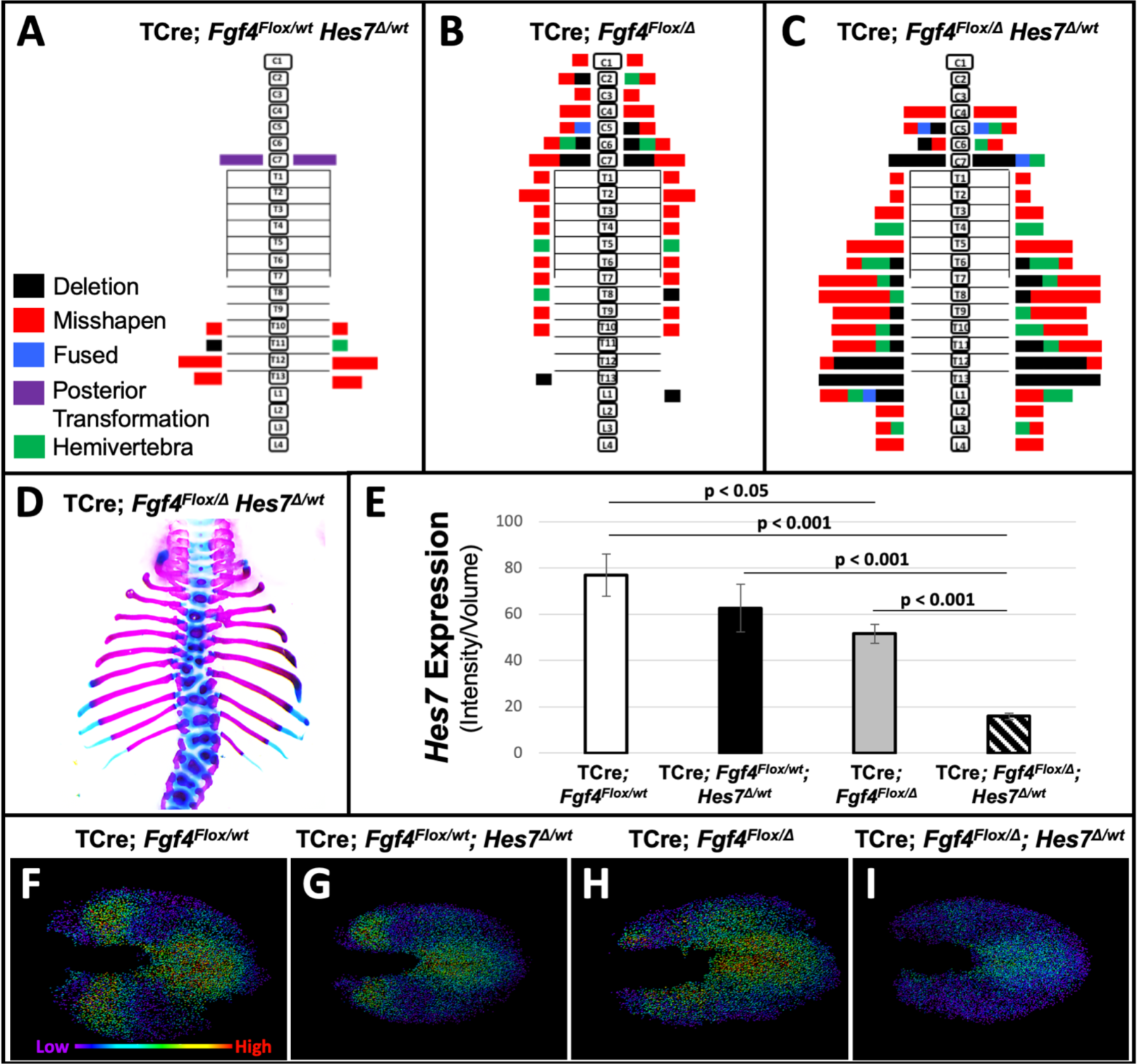
Removal of one *Hes7* allele exacerbates vertebral defects in *Fgf4* mutants. **A-C)** Histograms representing the vertebral column and associated ribs (C = cervical, T = thoracic, L = lumbar), showing variety and location of vertebral defects at E18.5 in the genotypes indicated (A and B, n = 7; C, n = 6). Each block in the histogram represents a single defect as indicated in the key in A. **D)** Skeletal preparation of a representative E18.5 compound *Fgf4* mutant-*Hes7* heterozygote (ventral view). **E)** Quantification of *Hes7* expression within the PSM in 5-6 somite stage embryos of the indicated genotype: control (n=10), *Fgf4*-*Hes7* double heterozygotes (n = 6), *Fgf4* mutants (n = 10), and compound *Fgf4* mutant-*Hes7* heterozygotes (n = 7). Quantification of *Hes7* mRNA expression, determined by measurement of fluorescent intensity per cubic micrometer within the volume of the mesoderm-specific *Tbx6* expression domain. Data in E are mean±s.e.m, significance determined by a Student’s t-test. **F-I)** Spot models based on HCR analysis of *Hes7* expression, in embryos of the indicated genotype, within the PSM, as defined by the *Tbx6* expression domain. Localized *Hes7* expression is colored by level of expression from low (purple) to high expression (red), as indicated. Images F-I are dorsal views, anterior left.

We performed a similar genetic analysis where we determined if *Hes7* heterozygosity likewise affects *Fgf8* mutant defects. We examined littermates of the last cross described in Table 1 (Figure 7 – figure supplement 1). *Fgf8* mutants in this cohort displayed the same homeotic transformations we previously observed with incomplete penetrance and expressivity: small ribs in the posterior cervical vertebra (3/5, 2 bilateral and 1 unilateral) or anterior lumbar vertebra (3/5, 1 bilateral and 2 unilateral). *Hes7* heterozygotes (also heterozygous for a floxed *Fgf8* allele) displayed a nearly identical frequency (2.7 defects per animal) as observed in the *Fgf4*/*Hes7* experiment (Figure 7A). Vertebral defects in compound *Fgf8/Hes7* mutants were not synergistic (4 defects per animal) but were clearly the *Hes7* heterozygous defects added to the relatively mild *Fgf8* defects (1.8 defects per animal, Figure 7 – figure supplement 1).

Therefore, we demonstrate that the relationship between *Hes7* and *Fgf4* is unique in that there is no genetic interaction between *Fgf8* and *Hes7*.

We then examined levels and spatial patterns of *Hes7* mRNA expression in each genotype from the *Fgf4/Hes7* mutant littermates, specifically within a volumetric model of the *Tbx6* expression domain, as we had in *Fgf4* mutants (Figure 5, 6). We observed a reduction in *Hes7* expression that correlated with the severity of vertebral defects within each genotype (Figure 7E). *Hes7* null heterozygotes had only a 19% reduction in expression, presumably because a loss of HES7 auto-repression results in enhanced transcription from the remaining wildtype allele. Such compensatory upregulation fails in compound *Fgf4/Hes7* mutants because we observed an 80% *Hes7* reduction to occur in these embryos (Figure 7E). Analysis of intensity-colored spot models of PSMs of these genotypes suggests a threshold effect of *Hes7* levels on oscillatory gene expression (Figure 7F-I). Oscillations appear relatively normal if *Hes7* levels are reduced 19% (in *Hes7* heterozygotes, Figure 7G) and disordered with a 33% reduction (in *Fgf4* mutants, Figure 7H). The synergistic 80% reduction of *Hes7* that occurs in compound *Fgf4/Hes7* mutants causes a severe dampening of oscillatory gene expression (Figure 7I). The resulting vertebral defects, though relatively severe (Figure 7A-D), are not as extreme as occurs in *Hes7* null homozygotes ^47^, indicating that this reduced level of *Hes7* supports limited patterning during segmentation. Together, our data support our hypothesis that FGF4 acts to maintain *Hes7* above a necessary threshold for normal somitogenesis. In *Fgf4* mutants, a reduction in *Hes7* levels, generates uncoordinated, asynchronous Notch oscillations. Disordered Notch oscillations initiate *Mesp2* unevenly in the anterior PSM leading to improperly shaped somites and subsequently, malformed vertebrae.

## Discussion

Here we describe mutants with only *Fgf4* or *Fgf8* inactivated specifically in the PSM. We found that *Fgf4* mutants display a range of cervical and thoracic vertebral defects that are caused by defective Notch oscillations during somitogenesis. In contrast, the vertebral columns of *Fgf8* mutants are normally segmented, with about 30% displaying minor homeotic transformations; such alternations in vertebral identity are not likely due to defects in somitogenesis *per se*.

Previously, a role for FGF signaling in wavefront activity in chick and zebrafish embryos was supported by pharmacological manipulation ^26, 50^ and in mouse mutants where both *Fgf4* and *Fgf8* are simultaneously inactivated using the same TCre activity that we use in this study ^28^. These double *Fgf4/Fgf8* mutants as well as mutants with tissue-specific inactivation of *Fgfr1* support a role for FGF signaling in clock oscillations, but these conclusions were complicated by a loss of PSM tissue due to potential wavefront defects ^16, 28, 31, 32^. Compared to these *Fgf* pathway mutants, the phenotype of *Fgf4* mutants is relatively subtle, with no wavefront defect, as indicated by no loss of caudal vertebrae ^42^ and no quantitative change in wavefront gene markers (Fig 2). This lack of any overt axis extension defect allows us to unambiguously identify *Fgf4* as an *Fgf* ligand gene required for a normal segmentation clock. This insight is supported by recently published *in vitro* work on the human segmentation clock ^51^.

Our mutants model human Segmentation Defects of the Vertebrae (SDV); *Fgf4* mutants resemble CS and the synergistic phenotype resulting from the additional removal of one *Hes7* copy in these mutants resembles human SCDO. Both of these diseases are caused by mutations in Notch signaling components that we find are misexpressed in *Fgf4* mutants, such as *HES7*, *MESP2,* and *LFNG* ^19, 20^. With regard to the human FGF pathway and SDV, a straightforward gene-disorder relationship has not been uncovered, probably because of the pleiotropic phenotypes of such putative mutations would preclude embryonic survival. Sparrow *et al* found that gestational hypoxia in mice results in an increase in the severity and penetrance of CS in *Notch1*, *Mesp2*, and *Hes7* heterozygotes ^25^. Intriguingly, CS is more severe in children living at high altitudes ^52^, suggesting a similar environmental effect may affect human development. In mice, hypoxia was found to diminish FGF signaling components, but expression of *Fgf8* was unchanged, and *Fgf4* was unexamined ^25^. Our data suggest that FGF4 may be part of the system that is responsive to this environmental insult. However, if this is the case, reduced FGF4 activity cannot be the only response to reduced oxygen levels because gestational hypoxia reduces expression of the FGF target gene *Spry4* in the PSM, whereas we found no change in expression in this or other such canonical FGF target genes in *Fgf4* mutants.

Rather, as is the case for hypoxia-treated embryos ^25^, *Hes7* expression is reduced in *Fgf4* mutants and we propose that this leads to aberrant segmentation. An indispensable tool to this insight was the use of multiplex HCR imaging ^33–35^. The simultaneous fluorescent imaging of multiple mRNA domains with this technique is particularly useful in a complex embryonic process, such as somitogenesis, where many complex and dynamic gene expression patterns need to be analyzed. HCR is a relatively new tool for mutant embryo analysis, just beginning to be used to analyze mutant embryos ^53–56^. We used HCR to examine the expression of multiple genes in a single tissue by combining it with Imaris image analysis software. This combination of HCR and Imaris modeling is broadly applicable for *in situ* quantification of gene expression within any complex embryonic and clinical sample. With distinct molecular markers, one can use this approach to quantify gene expression in any cell population with a precision that heretofore was not possible and still retain the intact whole mount embryo or tissue. Here, we generated and analyzed Imaris-based volumetric models of the PSM, based on *Tbx6* expression, and determined that *Hes7* levels are reduced by about 40-50% specifically in the PSM of *Fgf4*mutants during early somite stages when the progenitors of aberrant vertebrae are segmenting.

We propose that this reduction in *Hes7* expression is the primary defect in *Fgf4* mutants, and subsequently causes an irregular activated Notch (NICD) pattern, misexpression of *Mesp2*, and ultimately leads to vertebral defects. Given the molecular feedbacks during genetic oscillations in the PSM, we considered other possible models, but they do not fit our data. For example, we observed aberrant expression of *Lfng*, which encodes a glycosyltransferase that modulates Notch signaling ^57^. Although embryos completely lacking *Lfng* display aberrant *Hes7* oscillations ^32, 58^, overall *Hes7* mRNA levels are not reduced ^11^, unlike the case in our *Fgf4* mutants. Another factor required for *Hes7* transcription is encoded by *Tbx6* ^59, 60^, but we found no significant difference in expression of this gene. We support our proposal that the observed 50% reduction in *Hes7* is the cause of the *Fgf4* mutant with the synergistic worsening of the vertebral segmentation defect that occurs when we further reduce *Hes7* levels by removal of one wildtype allele. From another perspective, the mild CS caused by loss of one *Hes7* allele in a wildtype background ^25^ is worsened 9-fold by removing *Fgf4* (Figure 7). Therefore, *Fgf4* is a robustness-conferring gene that buffers somitogenesis against a perturbation in *Hes7* gene dosage ^61^.

We observed that in *Fgf4* mutants *Hes7* mRNA levels are reduced during rostral somitogenesis at E8.5 (5-6 somite stages), but are unaffected at about E9.5 (24-26 somite stages) when the embryo is beginning to generate correctly patterned somites that will differentiate into normally patterned vertebrae. Tam showed that between these two stages of mouse development, somites more than double in size ^62^. Such an increase can be due to a slower segmentation clock and/or faster regression of the determination front ^63^. Gomez et al. found that the caudal movement of the wavefront does not significantly change at these stages; therefore the clock must slow between E8.5 and E9.5 ^64^. Consistent with a slowing clock, we observed at least 2 and frequently 3 bands of *Hes7* expression at the *Fgf4*-sensitive stages (5-6 somite stages; Figure 4, 5), but only 1 to 2 bands when *Fgf4* loss has no effect on *Hes7* expression (24-26 somite stages; Figure 5 -figure supplement 5). Thus, it appears that FGF4 activity is required to maintain *Hes7* mRNA levels above a certain threshold when the segmentation clock is faster; when Notch oscillations slow, they may become FGF-independent, or other FGFs may be at play. If this is the case in all vertebrates, we might expect embryos with faster segmentation clocks, such as snakes ^64^, to have a longer window of FGF4-dependence for normal somitogenesis.

This study and past work indicate that FGF4 and FGF8 have both shared and unique roles in the vertebrate embryo’s posterior growth zone. They redundantly encode wavefront activities, preventing PSM differentiation ^28, 65^. Here, we show that FGF4 uniquely regulates Notch oscillations that control segmentation. Future efforts will explore why *Fgf8* loss-of-function causes the incompletely penetrant homeotic transformations that we observed - possibly FGF8 regulates *Hox* gene expression ^66^. These insights are intriguing in the context of the evolution of the vertebrate body plan. Vertebrates, together with Cephalochordates and Urochordates make up the phylum Chordata ^67^. Somitogenesis predates vertebrate evolution as it occurs in Cephalochordata, the most basal living chordates ^68^. In both Cephalochordates and Urochordates, the FGF essential for embryonic axis extension is an *Fgf8* ortholog ^69, 70^, suggesting that this *Fgf* (*“Fgf8/17/18”)* may be the ancestral gene in this process. However, *Fgf8/17/18* activity in these invertebrate chordates does not control gene oscillations as these embryos apparently lack a segmentation clock ^69, 71^. Therefore, we speculate that the recruitment of an *Fgf4* role during the evolution of vertebrate axis extension may have coincided with the development of gene oscillations that control somitogenesis.

## Materials and Methods

### Alleles, Breeding and Genotyping

All mice were kept on a mixed background. All genetic crosses are shown in Table 1, with the female genotype shown first and the male genotype shown second. PCR-genotyping for each allele was performed using the following primer combinations, *Fgf4flox*^72^ (5’-CAGACTGAGGCTGGACTTGAGG and 5’-CCTCTTGGGATCTCGATGCTGG), *Fgf4Δ*^72^ (5’-CTCAGGAACTCTGAGGTAGATGGGG and 5’-ATCGGATTCCACCTGCAGGTGC), *Fgf8flox*^73^ (5’-GGTCTTTCTTAGGGCTATCCAAC and 5’-GCTCACCTTGGCAATTAGCTTC) *Fgf8Δ*^73^ (5’-CCAGAGGTGGAGTCTCAGGTCC and 5’-GCACAACTAGAAGGCAGCTCCC), *Hes7Δ*^47^ (5’-

AGAAAGGGCAGGGAGAAGTGGGCGAGCCAC, and 5’-TTGGCTGCAGCCCGGGGGATCCACTAGTTC), TCre^36^ (5’-GGGACCCATTTTTCTCTTCC, and 5’-CCATGAGTGAACGAACCTGG)

### HCR

Hybridization chain reaction fluorescent in situs where carried out as described ^34^ with the modification of using 60 pmol of each hairpin per 0.5ml of amplification buffer. Hairpins where left 12-14 hours at room temperature for saturation of amplification to achieve highest levels of signal to noise^35^. Stained embryos were soaked in DAPI solution (0.5ug/mL DAPI in 5x SSC with 0.1% TritonX-100, 1% Tween20) overnight at room temperature. Split initiator probes (V3.0) were designed by Molecular Instruments, Inc.

### Whole Mount Immunohistochemistry, Colorimetric WISH, and Skeletal Staining

Notch Intracellular Domain staining (NICD) was performed following a significantly modified protocol^74^; briefly, embryos were dissected in cold PBS, briefly fixed for 5 minutes in 4% PFA, then fixed for 35 minutes in equal parts DMSO:Methanol:30% Hydrogen Peroxide at room temperature. They were then rinsed 3 x 5 minutes in 50mM Ammonium Chloride at room temperature, blocked in TS-PBS (PBS, 1% Triton X-100, 10% Fetal Calf Serum) for 30 minutes at room temperature, then incubated overnight in 1:100 anti-NICD (CST #4147) in TS-PBS at 4° C rocking. The next day, embryos were washed 4 x 10 minutes in TS-PBS, then incubated overnight in anti-rabbit-alexa647 (ThermoFisher A32733) 1:100 in TS-PBS at 4°C rocking.

Embryos were then washed 4x 10 minutes in TS-PBS then soaked overnight in DAPI, as described above in the HCR section, and embedded and cleared as described below. WISH and skeletal staining were performed as previously described ^75^.

### Embedding and Clearing

#### Embedding

Stained embryos where mounted in coverslip bottomed dishes suspended in ultra-low gelling temperature agarose (Sigma, A5030) that had been cooled to room temperature.

Once correct positioning of embryos was achieved the dishes were moved to ice to complete gelling.

#### Clearing

For tissue clearing we utilized Ce3D+, a modified version of Ce3D (Gerner et al., 2017), in which the concentration of iohexol (Nycodenz, AN1002424, Accurate Chemical & Scientific Corp) is increased to match the refractive index (RI) of the mounting solution to that of standard microscopy oils (nD = 1.515; not utilized in this study) and 1-thioglycerol is omitted to reduce toxic compounds within the solution (thereby making the solution safer to handle) and increase shelf life. To account for dilution by water from a sample (including agarose volume) we used Ce3D++, in which the iohexol concentration is further increased. Ce3D++ was designed to produce desired RI after two incubations - detailed protocol is available upon request. The protocol for preparing these solutions is as follows: in a 50 mL tube, add iohexol powder (20 g for Ce3D or 20.83 g for Ce3D++), then 10.5 g of 40% v/v solution of N-Methylacetamide (M26305, Sigma) in 1X PBS, and then 22.5 mg Triton-X-100 (T8787, Sigma) for a final concentration of 0.1%. The solution is mixed overnight at 37 C on an orbital rocker with intermittent vortexing then stored at room temperature. Embedded embryos were cleared using 2 changes of Ce3D++ solution while rocking at room temperature for a twenty-four-hour period.

#### Imaging

All images were obtained on an Olympus FV1000 confocal microscope, with an image size of 1620 x 1200 pixels and with a Kalman averaging of 3 frames. To capture the entire tissue a 10x UPlanApo objective (NA= 0.4) was used achieving a pixel size of 0.9um x 0.9um. Tissues were oriented in the same way with the anterior-posterior axis of the tissue oriented from left-right within the center of the field. Microscope settings were kept consistent between imaging.

Intensity calibration and shading correction was performed as previously described ^77^, however fluorophore dye concentrations used for shading correction were 0.05mg/L for fluorescein (Sigma 46960), 1mg/L for acid blue 9 (TCI CI42090), and 2mg/L for rose bengal (Sigma 198250). Dyes were placed in coverslip bottomed dishes. The Shading Correction plugin within Fiji was then used with the median flat field images acquired using the dye solutions.

#### Image processing

Images within figures were processed using Fiji ^78^ and represent max projections of z-stacks. Compared images are presented with identical intensity ranges for each channel. Orthogonal projections were made using Imaris software (Imaris V9.2.1, Bitplane Inc).

#### Statistical analysis

For all analysis at least 3 embryos were used unless otherwise stated in the text or caption. Significance was determined using a Students two-tailed t-test.

### Imaris Fluorescence Quantification and Modeling

#### HCR data

Flat-fielded image stacks were imported from Fiji into Imaris (Imaris V9.2.1, Bitplane Inc). A baseline subtraction was then performed for each probe, using a cutoff value specific for each probe and fluorophore combination. For all probes, except *Hes7*, the baseline cutoff was adjusted until signal from a tissue known not to express the gene was no longer detectable (e.g. neural epithelium for *Tbx6* ^40, 49^. For *Hes7*, this cutoff was determined by using embryos that were homozygous for a *Hes7* null allele ^47^ that lacked sequences complementary to the HCR probes; the cutoff was chosen that resulted in no signal in these mutants (Figure 5 - figure supplement 5).

The Surface model tool was used to build surfaces for each expression domain to be quantified except for *Fgf8* and *Hes7*, which were quantified using a surface derived from the *Tbx6* expression domain. Quantification of *Spry2*, *Spry4* and *Etv4* were determined by measurement of fluorescence intensity per cubic micrometer within the volume the respective gene expression domain. Surface models were generated using a surface detail value of 3uM, absolute intensity setting, and an absolute intensity threshold cutoff that was set to exclude background signal in tissues known not to express the gene being modeled. Once established for each probe, the same intensity threshold cutoff values were used for generating surfaces for all embryos. For generating *Tbx6* surfaces, a range of absolute intensity threshold cutoffs were used to achieve a final surface volume between 9.0e6 and 1.1e7 um3. Values for mRNA expression represent the intensity sum within the volumetric model divided by the volume of the model.

#### Immunostaining data

In images where NICD was immunostained, the surface creation tool was used. The mesoderm was selected, eliminating the ectodermal and endodermal tissue, on alternating z-planes through the entirety of the z-stack. The resulting pattern was interpolated for intervening planes. Values for NICD expression represent the intensity sum within the volumetric model divided by the volume of the model.

#### Spot modeling

Spot diameter was set to 4uM with a point spread function value of 8uM, local background subtraction was used, and sum intensity of the channel being modeled was used to threshold. Threshold cutoff values for spot models were set at 1,500 for all figures except Figure 6. In Figure 6 the thresholds values (a.u.) for the sum intensity are: low (blue), 1,000-6,000; low-medium (green), 6,000-11,000; medium (yellow), 11,000-16,000; medium-high (red), 16,000-21,000; high (white), 21,000 or greater. “Heat maps” in Figures 4 and 7 were generated within the Imaris software using a linear color scale; the scale in Figure 4 ranged from intensity values of 7,000 (blue) to 20,000 (red), the scale in Figure 7 ranged from 1,000 (blue) to 15,000 (red).

## Acknowledgements

We thank E. Kamiya and W. Heinz of the NCI Optical Microscopy and Image Analysis Lab, for their assistance with clearing using Ce3D+ and Imaris, respectively. We are grateful for technical assistance from T. Kuruppu, C. Elder, E. Truffer, and M. Boylan. We thank M. Kaltcheva, P. Abete-Luzi and A. Perantoni for critical reading of the manuscript.

---

Linked to Figure 1 is following figure supplement:

**Figure 1 - figure supplement 1.**
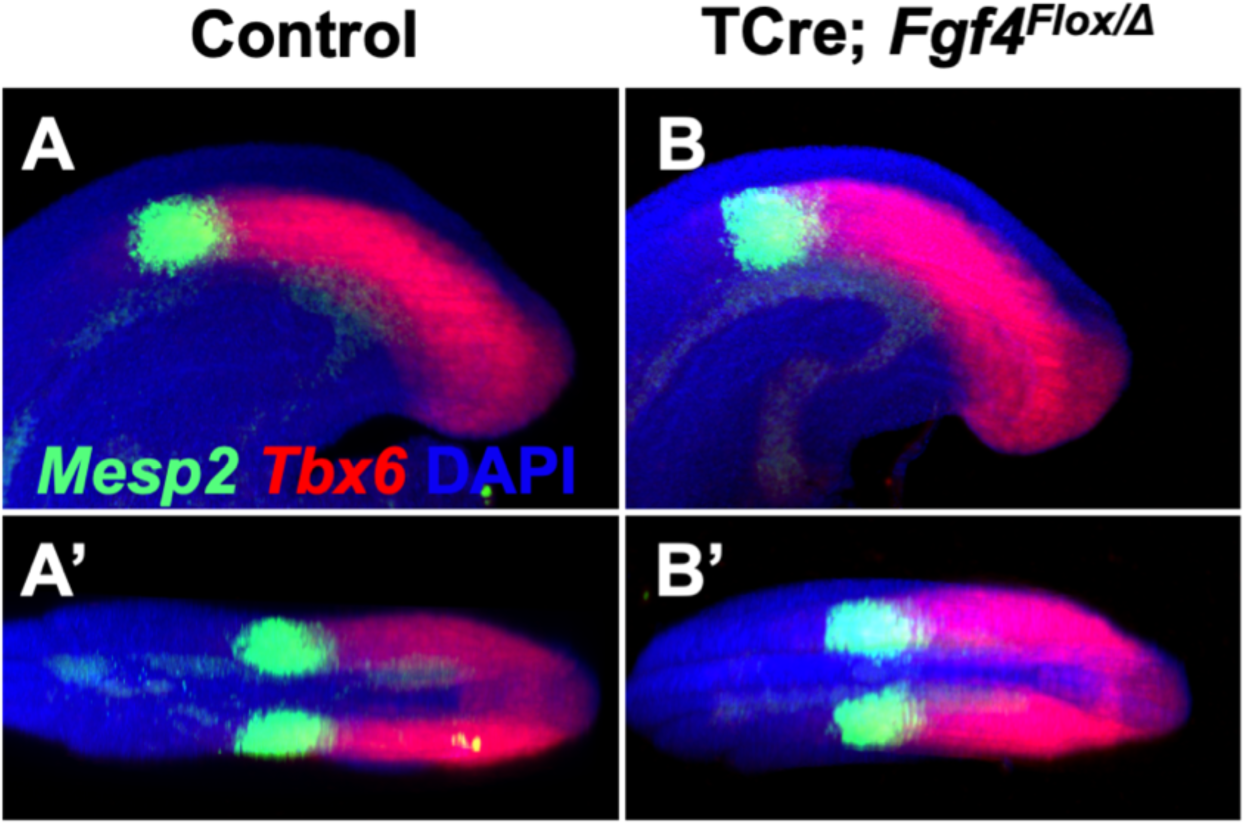
*Mesp2* pattern is normal at 24-26 somite stage in *Fgf4* mutants. **A-B’)** HCR staining showing no difference in the patterns of *Mesp2* and *Tbx6* expression between control and *Fgf4* mutant 24-26 somite stage embryos. A and B: MIP, lateral view with anterior to the left. A’ and B’: dorsal projections of A and B, anterior to the left. (control, n = 10; mutant, n = 11).

Linked to Figure 5 are the following figure supplements:

**Figure 5 - figure supplement 1.**
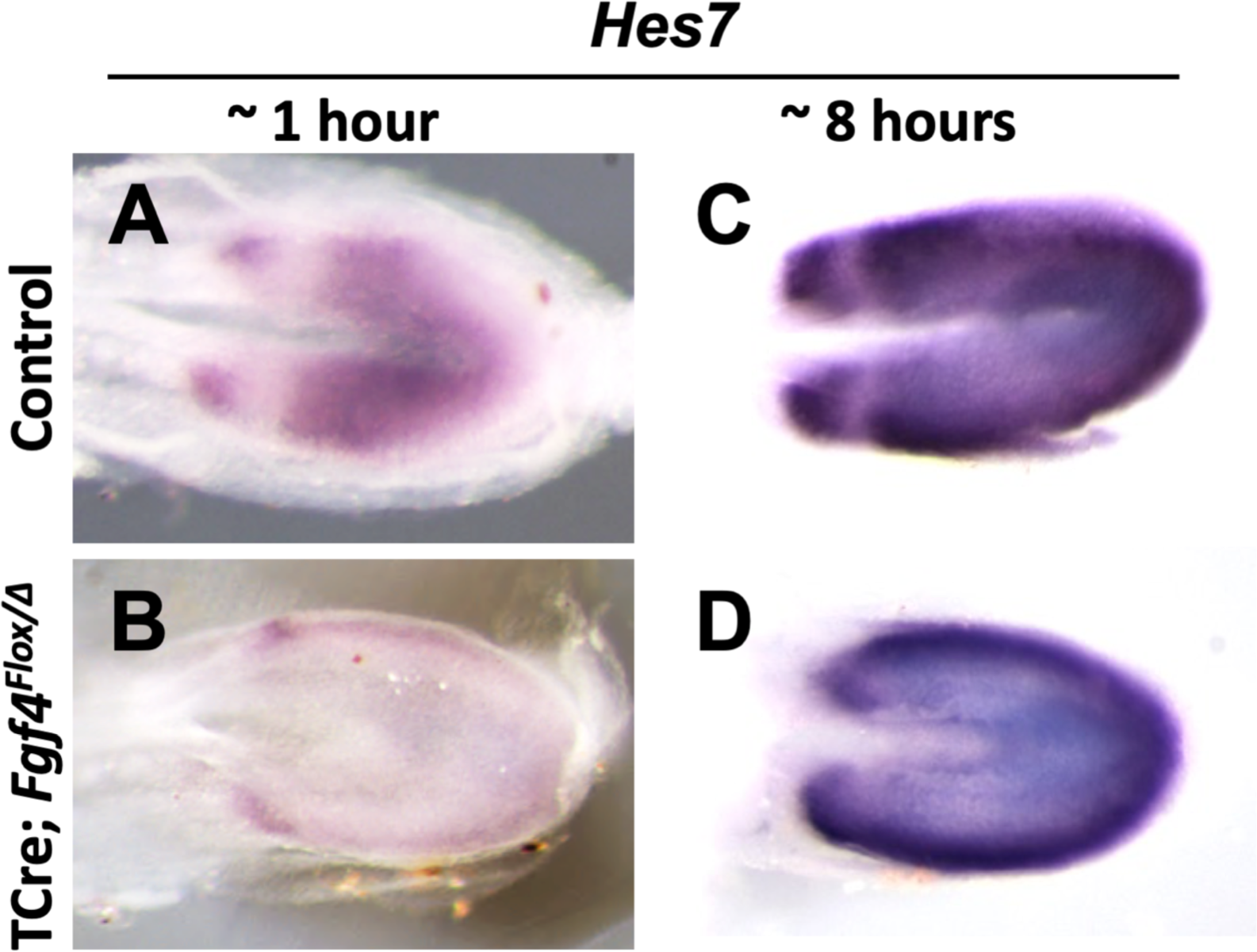
*Hes7* expression is abnormal in *Fgf4* mutants. **A-D)** Chromogenic whole mount in situ hybridization of *Hes7* mRNA in 5-6 somite stage control and *Fgf4* mutant embryos. Development of chromogenic signal was either not saturated (developed for ∼1hr in A, B) or saturated (developed for ∼8hr in C, D). Note decreased expression in *Fgf4* mutants (B) and abnormal pattern of oscillations (D). All images are dorsal views, anterior left and representative: A, n = 4; B, n = 4; C, n = 4; D, n = 6.

**Figure 5 - figure supplement 2.**
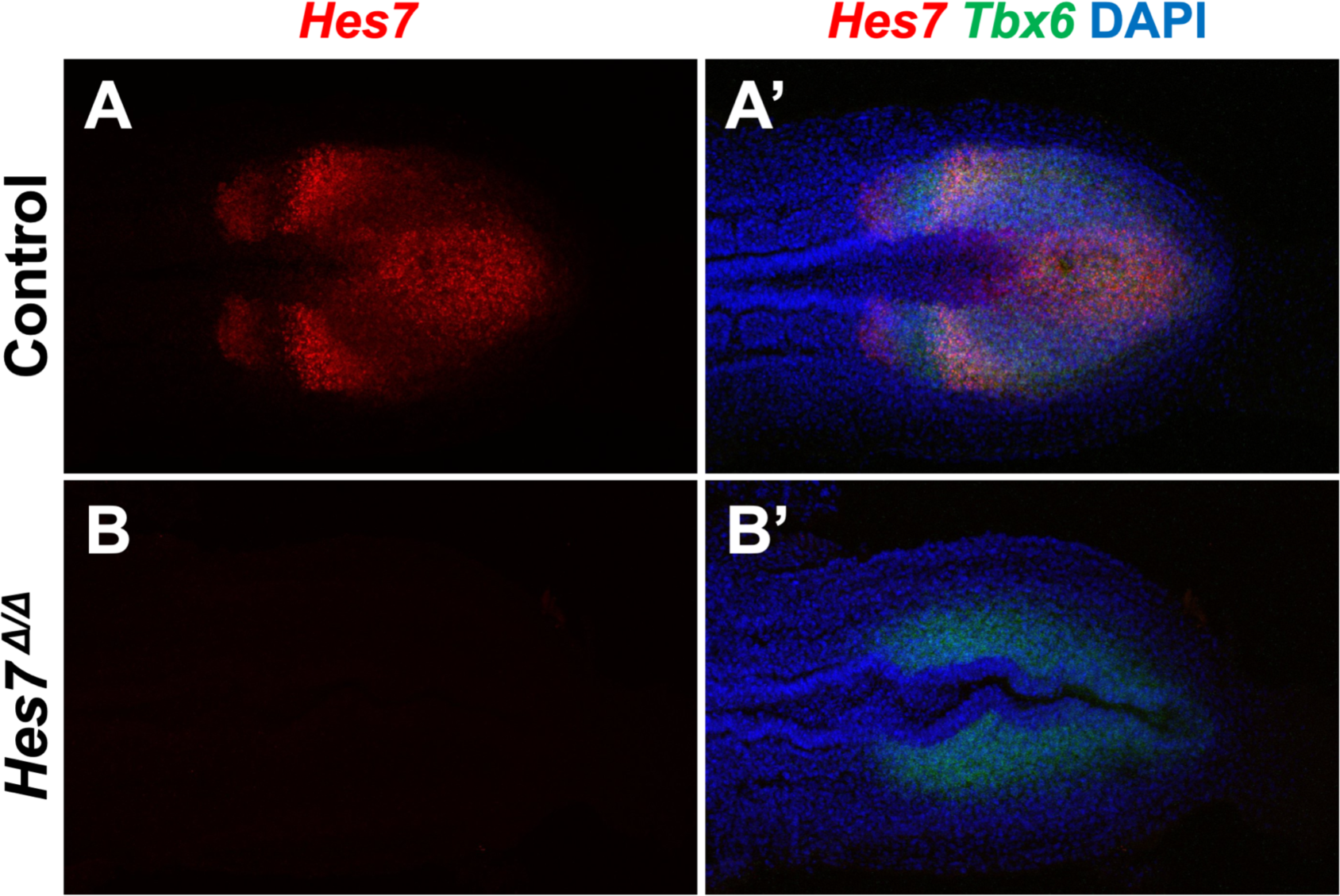
Using *Hes7* null homozygotes to adjust Imaris baseline cutoff. **A, B)** HCR staining of 5-6 somite stage control (n=3) and *Hes7* mutant (n=3) embryos using probes complimentary to the deleted region of *Hes7*. Note that the probe does not detect any product generated by the *Hes7* null allele. **A’, B’)** Composite image with *Tbx6* and DAPI staining in combination with *Hes7* of the same embryos as in A and B. All images are representative MIPs, dorsal view, anterior left. The baseline cutoff for Imaris-modeling of control and *Fgf4* mutant embryos was adjusted until a signal was absent from such *Hes7* null homozygotes. Same embryo shown in A, A’ and B, B’.

**Figure 5 - figure supplement 3.**
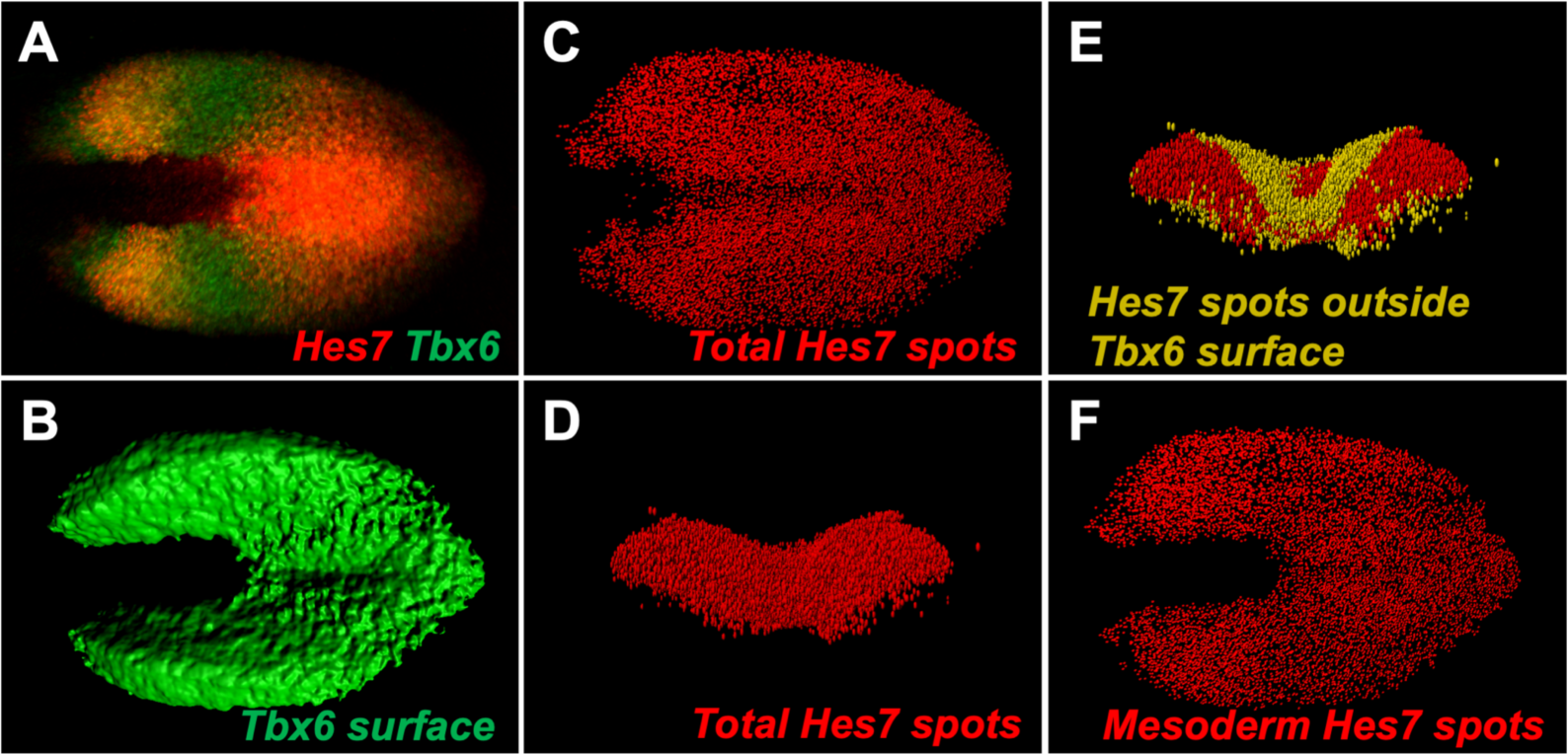
Imaris-modeling HCR Expression. **A)** HCR staining of *Hes7* and *Tbx6* in a 5-6 somite stage wildtype embryo; dorsal view, anterior left. **B)** Imaris-generated surface model of *Tbx6* expression domain from A. **C)** Imaris-generated spot modeling of *Hes7* expression domain from A. **D)** Anterior view of transverse projection of spot model in C. **E)** Spot model in D with *Hes7* spots outside of the *Tbx6* surface model colored yellow. **F)** Spot *Hes7* model from C with spots outside the *Tbx6* domain removed, thus only mesodermal spots remain. A similar *Tbx6* surface model was used in the quantification of *Hes7* expression levels in Figures 5H and Figure supplement 1 and 4. Spot modeling in F demonstrates how the spot models were generated in Figures 6 and 7.

**Figure 5 - figure supplement 4.**
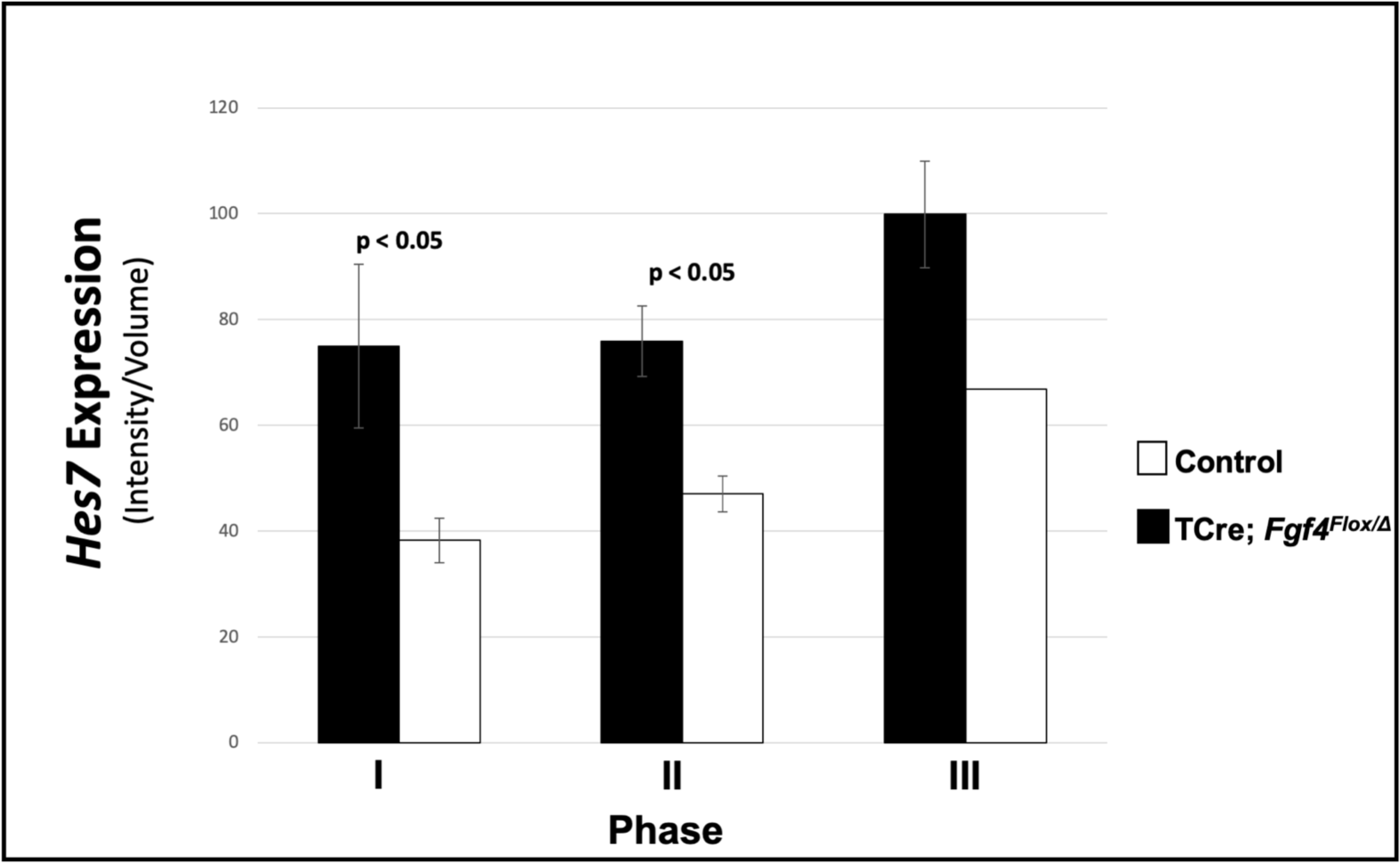
*Hes7* expression in *Fgf4* mutants is reduced, irrespective of oscillation phase. Quantification of *Hes7* expression within the PSM of control and *Fgf4* mutants at 5-6 somite stage. Data and embryos are identical to Figure 5H, except that fluorescence intensity quantification values are segregated by phase.

**Figure 5 – figure supplement 5.**
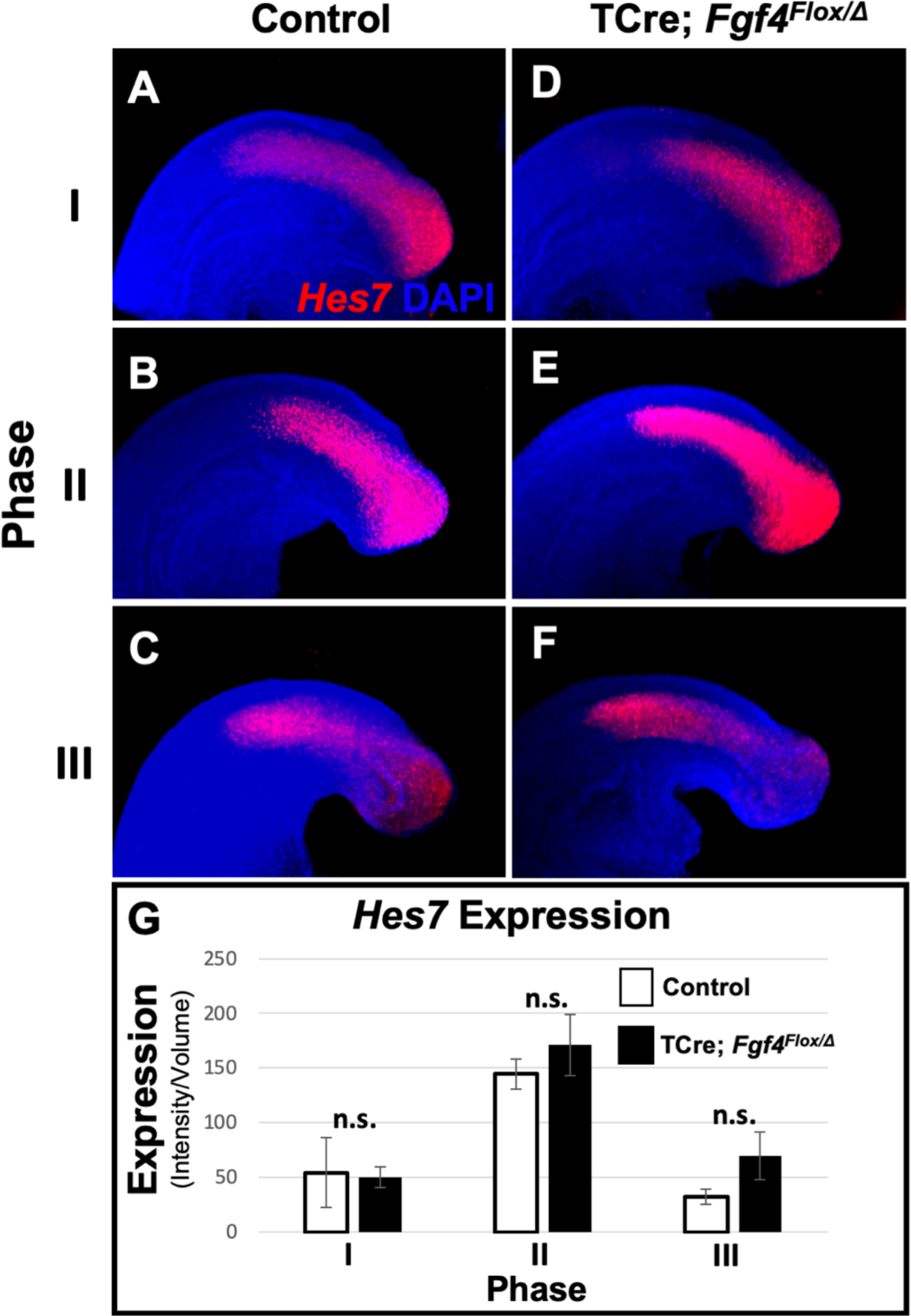
*Hes7* expression is normal at 24-26 somite stage in *Fgf4* mutants. **A-F)** HCR staining of representative 24-26 somite stage control and *Fgf4* mutant embryos showing normal patterns of *Hes7* expression for each phase of oscillation. MIP, lateral view with anterior to the left. **G)** Quantification of *Hes7* expression within the PSM, reveals no difference between controls and *Fgf4* mutants. Measurement of *Hes7* fluorescence intensity per cubic micrometer within the volume the mesoderm-specific *Tbx6* expression domain; data are mean ± s.e.m, significance determined by a Student’s t-test. Controls: phase I, n = 3; phase II, n = 3; phase III, n = 4. Mutants: phase I, n = 4; phase II, n = 4; phase III, n = 3.

Linked to Figure 7 is following figure supplement:

**Figure 7 - figure supplement 1.**
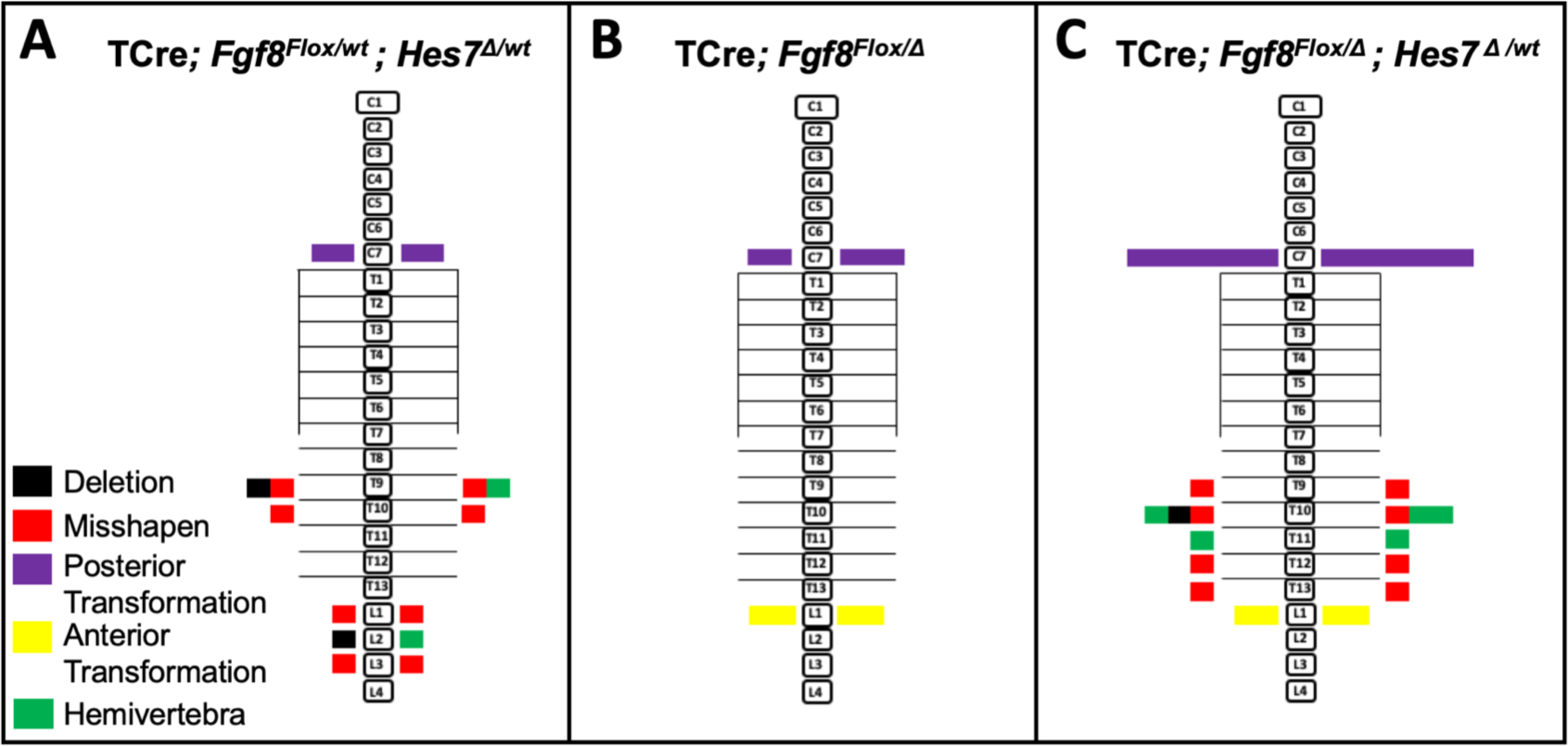
Removal of one *Hes7* allele has no effect on vertebral defects in *Fgf8* mutants. Histograms representing the vertebral column and associated ribs (C = cervical, L = thoracic, L = lumbar), showing variety and location of vertebral defects at E18.5 in the genotypes indicated (A, n = 6; B, n = 5; C, n = 8). Each block in the histogram represents a single defect as indicated in the key in A. Note that the vertebral defects in the *Fgf8*/*Hes7* mutants are additive, not synergistic.

